## Supplementary Figures 1-4 for "Sequences in the cytoplasmic tail of SARS-CoV-2 Spike facilitate expression at the cell surface and syncytia formation"

### Contents:

Supplementary Figures 1-4

Supplementary Tables 1 and 2 are provided separately as .xlsx files

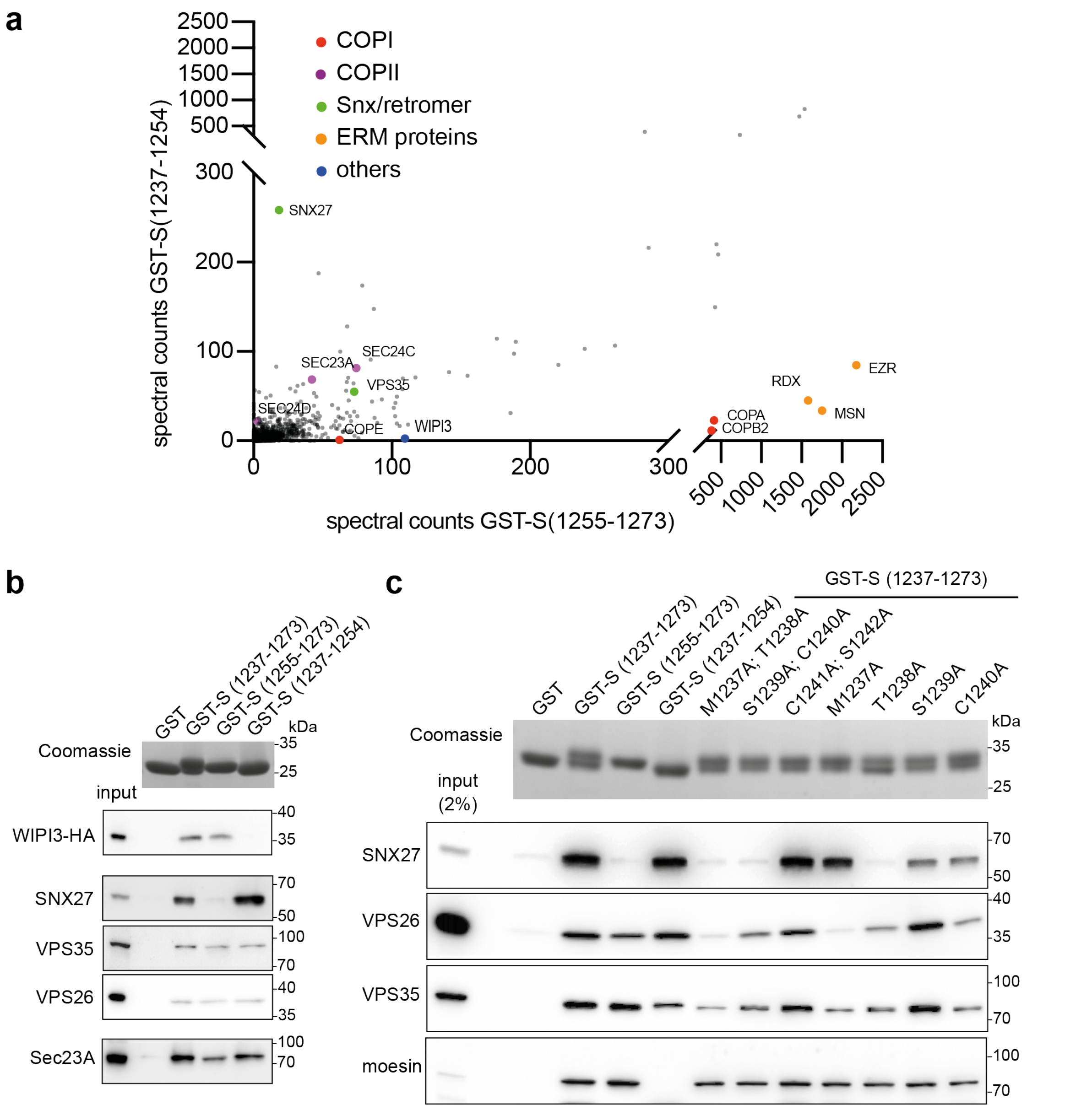


**Supplementary Fig. 1. Mapping of protein binding sites in the cytoplasmic tail of the SARS-CoV-2 S protein, including those of WIPI3 and SNX27/retromer**

**a** Plot comparing the spectral counts of proteins found by mass spectrometry analysis of interactors of membrane proximal GST-S(1237-1254) versus the membrane distal GST-S(1255-1273). The spectral counts are means from two biological repeats for GST-S(1237-1254) and three repeats for GST-S(1255-1273). All values in Supplementary Table 1.**b** The indicated GST-fusions expressed in E. coli, purified with glutathione Sepharose beads, and incubated with a 293T cell lysate followed by analysis of the bound proteins by immunoblot with the indicated antibodies. Coomassie-stained gel shows the GST-tail fusions. For WIPI3-HA, cells were transfected with a plasmid expressing the tagged protein and the blot probed for the HA tag. Representative blots from three independent experiments, input 1/50 of the lysate applied to the beads. **c** Mapping of residues required for binding to SNX27/retromer using the indicated GST-fusions as in (a) followed by probing with an anti-SNX27 antibody conjugated to HRP. Representative blots from three independent experiments, input 1/50 of lysate applied to beads.

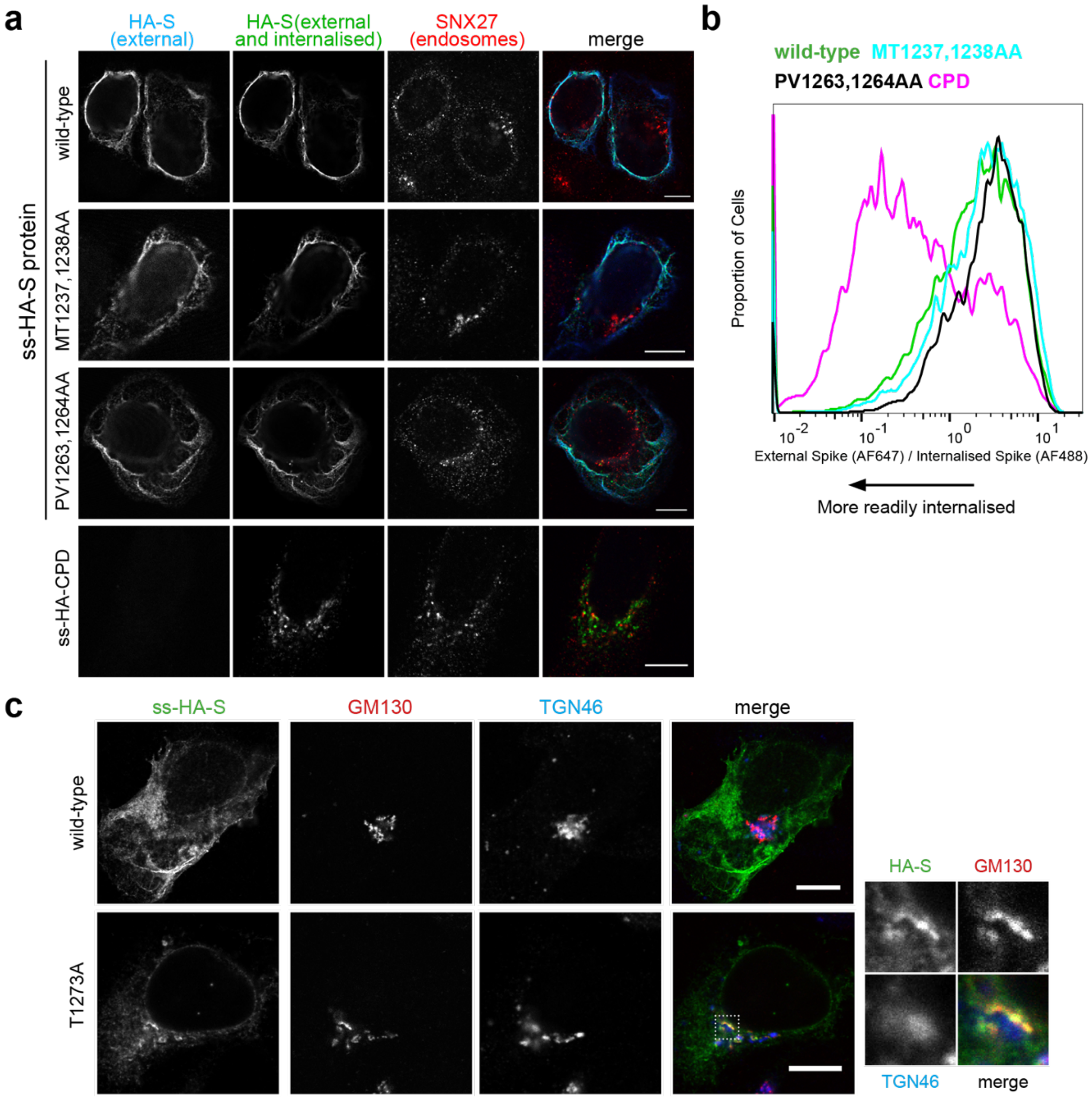


**Supplementary Fig. 2. The SARS-CoV-2 S protein is not rapidly internalised from the cell surface.**

**a** Micrographs of U2OS cells transiently expressing variants of S or carboxypeptidase D (CPD) (all with N-terminal HA tags), and subjected to an antibody uptake assay. Cells were incubated with an anti-HA antibody in complete medium for 30 minutes on ice, unbound antibody washed off and cells chased for one hour in complete medium at 37ºC. Cells were fixed and surface S was labelled using an AF647-conjugated secondary antibody under non-permeabilising conditions. Cells were then permeabilised and stained with an AF488-conjugated secondary antibody to label both surface and internalised S. Scale bars:10 µm. **b** Flow cytometry analysis comparing HA-tagged S and CPD in an antibody uptake assay. Cells were stained with an anti-HA AF488 conjugate for 30 minutes on ice, unbound antibody was washed off, and cells chased for 40 minutes at 37ºC. Cells were fixed and non-internalised anti-HA AF488 conjugated antibody was simultaneously quenched and relabelled with AF647 by incubation with an anti-AF488 antibody and an AF647-conjugated secondary antibody respectively under non-permeabilising conditions. Histograms represent the ratio of internalised S (AF488 signal) to that of non-internalised S (AF647 signal), and are normalised to the mode value and represent 5000 - 10,000 events, N = 2. **c** Confocal micrographs of U2OS cells transiently expressing N-terminally HA-tagged wild-type S or the T1273A variant, and labelled for GM130 (early-Golgi) and TGN46 (late-Golgi). Scale bars, 10 µm. Right panels show a magnified image of the Golgi region within the white dotted rectangle.

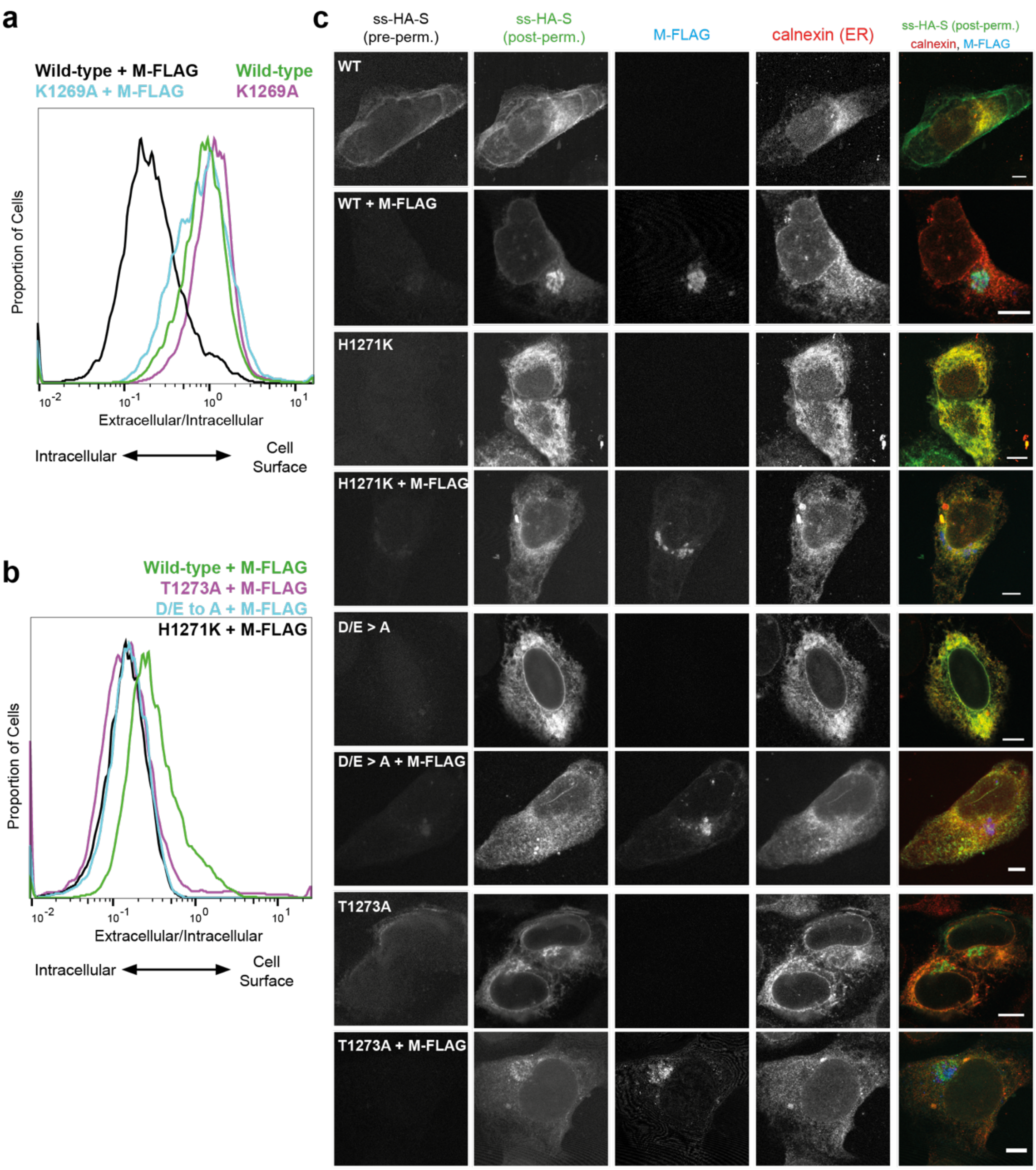


**Supplementary Fig. 3. Effect of co-expression of M protein on the localisation of S with mutations in the COPI and COPII binding motifs.**
**a** Histograms of the ratio of extracellular S (AF488 signal) to intracellular S (AF647 signal) for wild-type S and the K1269A mutant in the presence or absence of C-terminally FLAG-tagged M. As reported for SARS-CoV, Lys1269 of S is required for the protein to be retained in the Golgi by co-expressed M[1]. Histograms normalised to the mode value and represent ~10,000 events. Representative of three independent repeats. **b** As (**a**), except that the ratio of the extracellular and intracellular levels of wild-type S in the presence of M is compared to those of the indicated variants of S in the presence of M. Mutations that reduce COPII binding, or increase COPI binding, reduce further the proportion of S that reaches the surface in the presence of M. Histograms normalised to the mode value and represent ~10,000 events. Chi-squared tests show that in the presence of M, the differences between the median ratios of the wild-type (0.20) and of D/E>A (0.13), H1271K (0.11), and T1273A (0.12) are all statistically significant ($P<0.01$). Representative of three independent repeats. **c** Confocal micrographs of U2OS cells transiently expressing N-terminally HA-tagged S or variants, in the presence or absence of M-FLAG. Unpermeablised cells were stained for the HA tag on S, and then cells permeabilised and stained again for HA to label the internal pool of S, along with the FLAG tag on M, and the ER marker calnexin. Scale bars correspond to 10 µm.

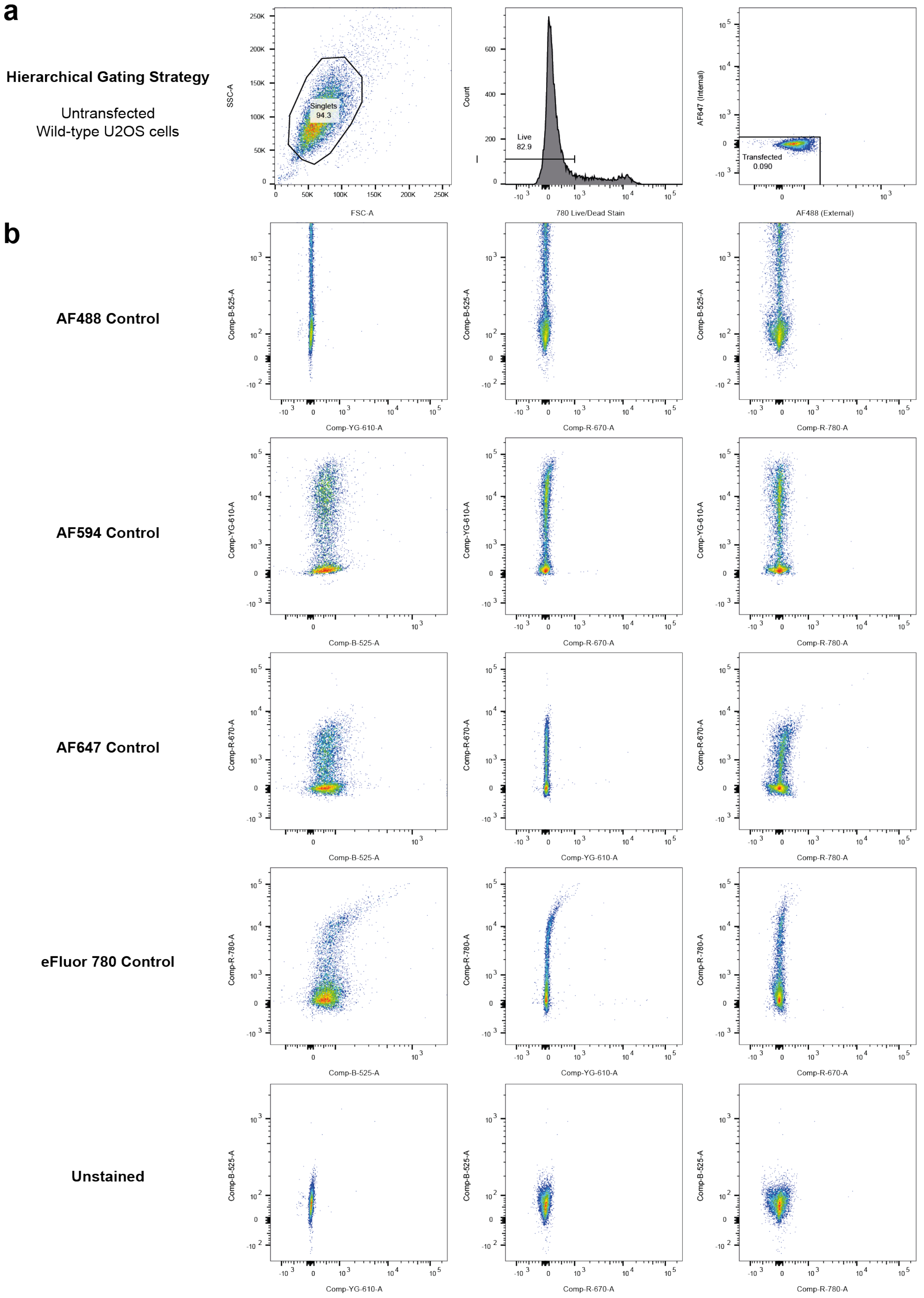


**Supplementary Fig. 4. Flow cytometry gating strategy and compensation controls.**
**a** Wild-type, unstained U2OS cells were used to define the hierarchical strategy to gate for singlet, live, transfected cells. The hierarchy is displayed in order from left to right. Transfected cells were defined as cells with an Alexa Fluor AF488 or AF647 signal greater

than the unstained control. Forward scatter (FCS) and side scatter (SSC) are also shown. **b** Cells stained with a single antibody or dye were used as single colour compensation controls. Where required for single colour epitope stains, cells were transfected with a plasmid expressing an epitope-tagged protein. Comp-B-525-A (AF488), Comp-YG-610-A (AF594), Comp-R-670-A (AF647), Comp-R-780-A (eFluor 780).
